# The CD74 inhibitor DRhQ improves cognition and mitochondrial function in 5xFAD mouse model of Aβ accumulation

**DOI:** 10.1101/2024.01.29.577832

**Authors:** Noah Gladen-Kolarsky, Cody J. Neff, Wyatt Hack, Mikah S. Brandes, Jack Wiedrick, Roberto Meza-Romero, Denesa R. Lockwood, Joseph F. Quinn, Halina Offner, Arthur A. Vandenbark, Nora E. Gray

## Abstract

Neuroinflammation and mitochondrial dysfunction are early events in Alzheimer’s disease (AD) and contribute to neurodegeneration and cognitive impairment. Evidence suggests that the inflammatory axis mediated by macrophage migration inhibitory factory (MIF) binding to its receptor, CD74, plays an important role in many central nervous system (CNS) disorders like AD. Our group has developed DRhQ, a novel CD74 binding construct that competitively inhibits MIF binding, blocks T-cell and macrophage activation and migration into the CNS, enhances anti-inflammatory microglia cell numbers and reduces pro-inflammatory gene expression. Here we evaluate its effects in β-amyloid (Aβ) overexpressing mice. 5xFAD mice and their wild type littermates were treated with DRhQ (100 µg) or vehicle for 4 weeks. DRhQ improved cognition and cortical mitochondrial function in both male and female 5xFAD mice. Aβ plaque burden in 5xFAD animals were not robustly impacted by DRhQ treatment nor was microglial activation, although in the hippocampus there was some evidence of a reduction in female 5xFAD mice. Future studies are needed to confirm this possible sex-dependent response on microglial activation as well as to optimize the dose, and timing of DRhQ treatment and gain a better understanding of its mechanism of action.

## 1. Introduction

Alzheimer’s disease (AD) is the most prevalent neurodegenerative disease in aging Americans [1] and is primarily characterized by deficits in cognition, especially memory loss and impaired executive function [2]. The highly complex and multifaceted nature of AD pathology renders the development of effective treatments beyond palliative care difficult [3]. While the accumulation of β-amyloid (Aβ) plaques and neurofibrillary tangles are the pathological hallmarks of AD, increased neuroinflammation and oxidative stress along with decreased mitochondrial function have been implicated in AD pathogenesis [4]. These processes are also interrelated with oxidative stress increasing the levels of proinflammatory cytokines that cause damage to mitochondria, which in turn results in even more oxidative stress within mitochondria [4, 5]. A similar bidirectional relationship exists in the AD brain between proinflammatory cytokines and Aβ plaques. Proinflammatory cytokines have been shown to increase the accumulation of Aβ plaques, while in turn the Aβ plaques themselves can also induce the release of proinflammatory cytokines [9]. The complex interplay between all of these biological functions has a compounding effect towards the progression of the AD, which underscores the importance of a multi-target approach for interventions [4].

Due to its relationship to so many essential cellular processes and because it is an early event in AD pathogenesis, modulating neuroinflammation has emerged as an attractive target for therapeutic intervention. Inflammation itself plays an important role in the context of fighting infectious disease; however, when it occurs chronically as in AD, inflammation can accelerate disease progression [6]. The innate immunological process of neuroinflammation is mediated by multiple cell types and secreted factors. In the CNS, microglia are the resident immune cells. In a healthy brain, microglia play an important role in synaptic pruning and elimination of damaged or apoptotic neurons via efferocytosis [7]. Microglia also mediate the phagocytosis of misfolded proteins, such as aggregated Aβ [8]. Under normal homeostatic conditions the activity of microglia is inhibited by dedicated pathways that suppress unwanted inflammatory responses [9]. However, in a disease state like AD, activated microglia can engulf neuronal synapses and secrete inflammatory factors that can injure neurons, both directly and indirectly [10].

Microglial activation and neuroinflammation are evident during the preclinical phase of AD [11] and elevated inflammatory markers are seen in both blood and the brains of AD patients [12]. One such elevated inflammatory marker is macrophage migration inhibitory factor (MIF). Elevated MIF expression has been observed in the plasma and CNS of AD patients [13, 14]. MIF-induced signaling through its cellular receptor, CD74, represents a common inflammatory pathway for many neurodegenerative conditions including AD [15] and in the AD brain, MIF-related increases in CD74 have been observed in Aβ plaques and [16] neurofibrillary tangles as well as microglia [17]. These changes in MIF levels and signaling have been recapitulated in rodent models of AD [18, 19].

Members of our group have developed a class of compounds that competitively inhibit MIF binding to CD74 and show promise as therapeutic agents in neurodegenerative disease [20]. The partial major histocompatibility complex (pMHC) class II construct, DRα1-MOG-35-55 (DRα1) has been shown to reduce disease severity and attenuate inflammation in the experimental autoimmune encephalomyelitis (EAE) mouse model of multiple sclerosis through direct inhibition of MIF binding to CD74 [16] [21]. Similar beneficial effects on disease severity and inflammation were also seen in models of stroke, traumatic brain injury, and methamphetamine addiction [20, 22–25]. A more recently developed iteration of this construct, with single amino acid substitution (L50Q) called DRhQ was shown to have 8-10 fold higher affinity for CD74 [26], suggesting it could have an even more potent neuroprotective effect on MIF-CD74 relation inflammation.

In this study, we investigated the effects of disrupting MIF-CD74 signaling in the context of AD. Using the 5xFAD mouse model of Aβ accumulation we evaluated the effects of DRhQ treatment on cognition, neuroinflammation, mitochondrial function and antioxidant response.

## 2. Methods

### 2.1 DRhQ synthesis

The DR⍺1 constructs purification has been previously reported [14]. A DNA fragment containing the sequence for MOG-35-55 peptide, a flexible linker and the MHC Class II DR⍺1 domain from amino acid residues 15 through 97 and a similar synthetic DNA fragment containing the mutation L50Q in the DR⍺1 domain were cloned. These clones were transformed into the E. coli strain BL21, plated onto agar plates, then subsequently, cultured, grown and harvested by centrifugation. Finally, following resuspension and sonication, protein collected from inclusion bodies were solubilized and purified. Fractions were collected and analyzed using SDS-PAGE, and fractions containing the protein of interest were pooled together, concentrated and flash frozen and stored until ready for use [13].

### 2.2. Animals

All experiments were conducted in accordance with the NIH Guidelines for the care and Use of Laboratory Animals and were approved by the institutional Animal Care and Use Committee of the Veterans Administration Portland Health Care System (VAPORHCS; IACUC #4688-21). 5xFAD and B6SJLF mice were purchased from Jackson Laboratory, Bar Harbor, ME. Animals were housed in a climate-controlled facility with a 12h light/dark cycle. Animals were fed PicoLab Laboratory Rodent Diet 5LOB (Lab Diet, MO) and provided with water and diet ad libitum. The mouse colony was maintained by breeding 5xFAD positive male mice with B6SJLF1 females. Wild type (WT) littermates were used as controls for each experiment.

At six months of age, male and female 5xFAD mice and WT littermates began treatment with either 0 or 100ug DRHQ in 20mM Tris administered subcutaneously five times per week for the first week followed by three times per week in the following three weeks for a total of 4 weeks of treatment (Figure 1a). In the finals ten days of treatment mice underwent cognitive testing. The 61 mice in the experiment (Figure 1b) were tested in two cohorts. Although the cohorts contained some imbalances each included male and female mice of both genotypes (5xFAD and WT) and treatment conditions (vehicle and DRhQ) with at least two from each group represented in each cohort.

**Figure 1:**
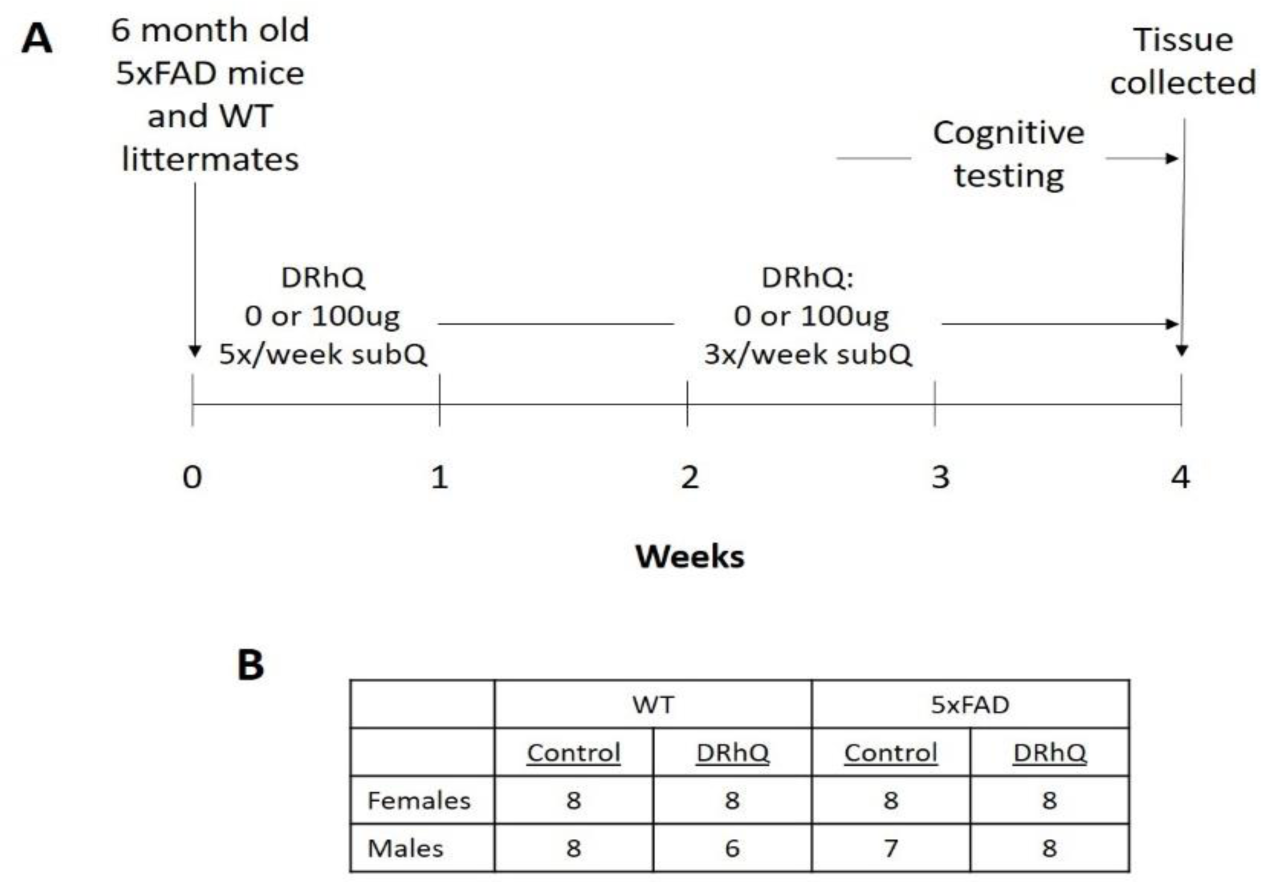
Experimental design. A) Treatment timeline and B) number of animals in each treatment group.

### 2.3 Cognitive analyses

#### •2.3.1 Novel Object Recognition Test (NORT)

The NORT assessment of recognition memory was performed as previously described [15]. Briefly, a rectangular open field apparatus (39 cm x 39 cm x 39 cm) is used for the experiment. In the habituation phase, mice were placed in the apparatus for 5 minutes without any objects placed in the field once per day for two days. On the third day mice underwent the training phase, where they were introduced to the arena now containing two identical objects and allowed to explore for three separate 10-minute training sessions each one hour apart. Testing followed 2 hours and 24 hours after the conclusion of the final training session, where mice were again placed in the apparatus but this time one of the objects from training was replaced with a novel object. The novel object was distinct for the 2 hour and the 24-hour tests. Mice were allowed to explore the two objects for 5 minutes during each test. Time spent exploring the novel object relative to the total time exploring both objects was quantified as a percentage. The objects used were a glass cylindrical votive, a metal cylindrical jar, a plastic rectangular box, and a plastic trapezoidal prism, all with approximately the same height. The object used and whether it was novel or familiar was balanced between groups. No preference was observed in this study for any object over the others.

#### 2.3.2. Conditioned Fear Response (CFR)

The CFR test of contextual memory was performed, as previously described [16], after NORT following a two-day rest period. CFR occurred in three phases: habituation, conditioning and testing. In the habituation phase, animals were allowed to freely explore a 16×16×12 inch chamber with a wire floor for 5 minutes during which baseline freezing is measured using ANY-MAZE software (Stoeling Co., Wood Dale, IL, USA). The conditioning phase followed immediately after habituation, where mice were exposed to three random 1-second shocks (0.5mA) using the Fusion v6.0 for Superflex Edition Software (Omnitech electronics, Inc., Columbus, OH, USA). 24 hours after conditioning, mice were reintroduced to the same chamber and freezing was quantified over two 5 minute periods in the absence of any additional shocks. Freezing time for each animal was represented as the difference in the freezing time observed during testing versus during the habituation period to account for any baseline differences in overall activity.

### 2.4. Euthanization and tissue collection

Following behavioral tests, mice were euthanized using inhalable isoflurane and cerebral dislocation, and brain samples were collected. The anterior 3mm of bilateral prefrontal cortex was dissected and frozen. The right hemisphere of each sample was immersion-fixed in 4% paraformaldehyde (PFA; Fisher Scientific, MA), diluted in 10x phosphate buffered saline for immunohistochemical analysis. The contralateral cortex was subdissected and divided in half. One half was used for gene expression analysis, and the other for synaptosomal isolation and analysis of mitochondrial function.

### 2.5. Immunohistochemistry (IHC)

Right hemispheres were incubated in 4% PFA for 24 hours at room temperature. Following PFA incubation, the tissue was transferred to 1x phosphate buffered saline (PBS) and incubated for a further 24 hours, then subsequently transferred to 15% and 30% sucrose solutions for 24 hours each before being frozen at -80°C for sectioning. Frozen coronal sections (40uM) were embedded in Optimal Cutting Temperature (O.C.T.) Compound (Sakura Finetek USA, Inc., Torrance, CA, USA) and cut on a freezing microtome. Sections were then stored in a sectioning solution (15% glycerol, 10% TBS, diluted in diH2O) before subsequent processing.

Similar depth sections were placed in a quenching solution (30% methanol, 10% hydrogen peroxide, 10% Tris-HCl buffered saline; TBS) for endogenous catalase activity and then blocked (2% bovine serum albumin, 10% horse serum, 2% triton x, 10% 10x TBS, diluted in diH2O). Sections were then incubated with one of the two following primary antibodies validated in our previous work [27], diluted 1:1000 in PBS: anti-AꞴ polyclonal antibody (Thermo Scientific, MA #44-136); biotinylated *Griffonia simplicifolia* Lectin I (GSL 1, Vector Labs, CA #B-1205).

For Aβ, biotin-labeled anti-rabbit secondary antibody (1:200, Invitrogen, REF 65-6140, LOT WA314056), and an avidin-linked peroxidase complex (AB-HRP kit #PK-4000, Vector Laboratories) were used to visualize staining. DAB (Sigma Fast 3,3 Di-aminobenzidine Tablet Set, D-4418 (Sigma-Aldrich Corp, St. Louis, MO, USA)) was used as a counterstain as previously described (Harris et al. 2014). For biotinylated GSL staining, samples were moved directly to avidin-linked peroxidase complex for visualization. DAB was subsequently used as a counterstain.

Sections were then mounted on UltraSlips coverglass slides (GC2450-ACS) and scanned using PrimeHisto XE (Pacific Image Electronics, Torrance CA, USA). ImageJ software (Rasband, W.S., ImageJ, U.S. National Institutes of Health, Bethesda, Maryland, USA, https://imagej.nih.gov/ij/, 1997-2018) was used to quantify images. Images were converted to 8-bit grayscale, and a human use employed the polygon tool to encompass the hippocampus or cortex and the total area of each brain area was recorded. The threshold was adjusted to remove ubiquitous background staining, highlighting only areas of intense staining. Staining was quantified in three coronal sections representing regions of anterior, middle and posterior hippocampus and cortex from each right hemisphere sample. Hippocampal and cortical areas were traced using a computerized stage and stereo investigator software. The extent of pan-AꞴ pathology and GSL staining was expressed as a percentage of the cortical or hippocampal area (cm2) that was occupied by detectable immunoreactive staining. Mean values for each parameter were calculated using at least three sections per animal.

### 2.6. Gene expression analysis

One half of the cortex from the left hemisphere of each mouse brain was frozen at -80°C. RNA was extracted from homogenized deep gray tissue with a QIAsymphony RNA kit (QIAGEN) using the RNA CT 400 v7 protocol. Reverse transcription was performed using a SuperScript™ VILO™ cDNA synthesis kit (Invitrogen). Relative gene expression was determined using custom Taqman® Gene Expression Array Cards (Thermo Fisher Scientific) pre-loaded with the following commercially available Taqman primers: interleukin-6 (IL6), tumor necrosis factor alpha (TNFα), C-chemokine receptor type 6 (CCR6), CD36, CD74, C-X3-C motif chemokine ligand 1 (CX3CL1), C-X3-C motif chemokine receptor 1 (CX3CR1), Advanced Glycosylation End-Product Specific Receptor (AGER or RAGE), triggering receptor expressed on myeloid cells 2 (TREM2), complement C3a receptor 1 (C3AR1), mitochondrial encoded NADH dehydrogenase 1 (mt-ND1), mitochondrial encoded cytochrome c oxydase 1 (mt-CO1) and mitochondrial encoded ATP synthase F0 subunit 6 (mt-ATP6) and glyceraldehyde-3-phosphate dehydrogenase (GAPDH). Quantitative PCR (qPCR) was performed using a QuantStudio™ 12K Flex Real-time PCR System (Applied Biosystems) and a QuantStudio™ 7K Flex Real-time PCR System (Applied Biosystems). Samples were run in triplicate across 29 custom array cards and data was normalized to GAPDH using the delta-Ct method, oriented so that larger values indicate higher expression.

### 2.7. Analysis of mitochondrial function

Mitochondrial bioenergetics were quantified in cortical synaptosomes using the Seahorse Xfe96 Analyzer. Synaptosomes were freshly isolated from one half of the cortical tissue from the left hemisphere using Syn-per reagent (Thermo Scientific #87793) as per the manufacturer’s protocol. The total protein concentration of the synaptosomal preparation for each mouse was determined using a bicinchoninic acid (BCA) assay. A total of 10ug of synaptosomal protein diluted in 25uL mitochondrial assay solution (MAS; 70mM sucrose, 220mM mannitol, 10mM KH2PO4, 5mM MgCl2, 2mM HEPES, 1mM EGTA, 0.2% BSA) was plated in each well of a polyethylenimine (PEI) coated 96-well Seahorse plate (5-6 replicate wells for each animal) and the plate was centrifuged at 1200g for 1h at 4C. 155 μL of Agilent Seahorse XF DMEM Medium pH 7.4 (Part # 103575-100) supplemented with 1 mM pyruvate, 2 mM glutamine, and 10 mM glucose was plated in each well of the 96-well plate.

Mitochondrial function was then assessed using the MitoStress kit (Agilent #103015-100) as previously described [28]. Briefly, oxygen consumption rate (OCR) was measured under varying conditions. After three initial baseline measurements of OCR, the ATP synthase inhibitor oligomycin (2μM) was added and three subsequent measurements were taken. Next an electron transport chain (ETC) accelerator, p-trifluoromethoxy carbonyl cyanide phenyl hydrazone (FCCP at 2 μM), was added and three measurements of maximal respiration were taken. Finally, the mitochondrial inhibitors rotenone (0.5μM) and antimycin (0.5μM) were added and three final measurements were taken. ATP-linked respiration was calculated by subtracting the average of the three time points after administration of oligomycin from the average of the three basal times points. Spare capacity was calculated by subtracting the average of the basal measurements from the average of the three maximal measurements following FCCP administration.

### 2.8. Data normalization

Since the mice were tested in two cohorts, systematic batch differences arose for some of the sets of measurements that required normalization across the two cohorts prior to statistical analysis. In addition, there were small differences in the joint distribution of sex and age in the two cohorts of mice (for logistical reasons it was not possible to exactly balance the cohorts with respect to these factors as would be ideal), and these differences were sometimes influential on measurements, necessitating regression adjustment. In order to decide on an appropriate set of normalization adjustments for each set of data in an unbiased way, we fit a regularized (LASSO) linear regression model to each outcome, including fully factorial specification of the experimental factors (genotype, sex, treatment and all their two- and three-way interactions) as forced model terms, and as unforced terms we additionally added age at sacrifice and testing cohort (1 or 2) along with their interaction and further two-way interactions of each factor with genotype, sex, and treatment. We used the minimum-BIC (Bayesian Information Criterion) rule to determine the regularization parameter and examined the resulting model for indications of potentially nonlinear batch effects due to cohort membership or cohort imbalances in age or sex distribution; if the minimum-BIC LASSO included unforced terms (rather than setting their coefficients to zero) then we performed normalization adjustments appropriate to the terms included. The motivation for this search is that in the presence of batch confounding, measurements may be systematically higher in one batch than the other (regardless of treatment or other factors), or measurements may show a trend over age at sacrifice (perhaps independently of batch), or effects of other factors may vary across specific combinations of batch and age due to simultaneous imbalance in age and some other experimental factor across batches. Data normalization adjustments indicated by the minimum-BIC LASSO were implemented outcome by outcome using separate regression models, with the caveat that we required substantially similar outcomes (e.g. NORT at 2 hours and NORT at 24 hours) to be normalized in the same way. Briefly, the adjustments were: 1) for NORT outcomes, no adjustments needed; 2) for CFR outcomes, adjustment for main effect of cohort only, using absorbing linear regression; 3) for IHC outcomes, due to very complex confounding of nuisance age associations with testing cohort (batch effects complicated by age associations observed over differing ranges of age in each cohort), we first performed an alignment of age associations between cohorts for GSL and AꞴ simultaneously using a generalized (log-gamma) structural equation model where the cohort adjustments to the age associations were forced to be the same for each type of outcome in each region (i.e. the cortical and hippocampal GSL measurements shared a common adjustment and the cortical and hippocampal AꞴ measurements shared a common adjustment), followed by age-normalization of all the measures by subtracting the age trend from linear regression; 4) for the gene expression data (which was already normalized by GAPDH expression), no adjustments needed; and 5) for the mitochondrial function outcomes, a common adjustment for main effects of both age and cohort was enforced by calculating an adjustment factor for the geometric mean of the basal, maximum, and ATP-linked consumption values using residualized predictions from a log-gamma regression model. For outcomes where normalization was performed, the normalized values were used in all subsequent statistical analyses.

### 2.9. Statistical analysis

For all outcomes in each testing domain (NORT 2h and 24h; CFR 0-5m and 5-10m; immunohistochemistry of GSL and AꞴ plaque burden; expression of mitochondrial and inflammatory genes; and mitochondrial function measured as rates of basal, maximum, spare, and ATP-linked O_2_ consumption), we fit a regression model of the outcome as a function of a fully factorial specification of the experimental design factors (genotype, sex, and treatment) that includes all possible interactions, but the functional form of the regression model differed depending on the metric and distributional features of the outcome. The model forms utilized were: 1) fractional logistic regression for both NORT outcomes; 2) ordinary linear regression for both CFR outcomes; and 3) log-gamma regression (a kind of nonlinear regression expressed as a generalized linear model that naturally accommodates skew in the outcome distribution) for all gene expression, IHC, and mitochondrial function outcomes. Additionally, for the two NORT outcomes we also evaluated the rates of participation (i.e. whether or not the mouse investigated any objects at all during the test) using a linear probability model to check for differences between testing cohorts, genotype, sex, or drug treatment groups. Leave-one-out jackknife estimation of standard errors was employed in all models because it is nearly unbiased for smooth statistics (such as regression coefficients) even at small sample sizes and robust to violations of distributional assumptions in the model. Group comparisons were implemented using model contrasts of predictive margins calculated under the assumption of balanced groups (i.e. all group margins are given equal weight in forming the contrast), and all contrasts are annotated with 95% confidence intervals or p-values as appropriate. Note that we made no adjustments for multiple comparisons, and since this is an exploratory study we imposed no threshold for significance; instead, all p-values are presented as calculated and any judgments about presence of effects are presented qualitatively rather than deterministically. Statistical analyses and figures were implemented using Stata v18 (StataCorp, College Station, TX, USA).

## 3. Results

Missingness was assessed for all endpoints and no salient or systematic effects were observed regardless of treatment group or cohort (Supplementary Table 1). All results below are represented graphically as the change between vehicle (blue diamond) and DRhQ treatment (yellow diamond). The top portion of each graph shows the overall DRhQ treatment effect for a given genotype with data from male and female animals combined together (green line = WT and orange line = 5xFAD; slope and confidence interval (in brackets) are also listed) with the diamonds indicating group averages. Confidence intervals that do not span 0 indicate a difference that excludes the null value with 95% confidence at this sample size. Individual points for each animal are plotted below, separated out by sex and again with diamonds indicating group averages. The p-values listed refer to the compatibility of the change with the null value in the DRhQ-treated group relative to the vehicle-treated group for a given sex (pink line = female, blue line = male).

### 3.1 DRhQ improves recognition memory but does not appear to affect associative memory in 5xFAD mice

In the 2h NORT test there was a pronounced deficit in performance in the 5xFAD vehicle treated mice as compared to the WT vehicle treated mice (p<0.001). DRhQ improved NORT performance in male and female 5xFAD animals during the 2h test (Figure 2A). There was no apparent interaction between sex and treatment effect in the 5xFAD mice (p=0.722). In the 24h test, the genotype effect in the vehicle treated animals was similar to what was seen at 2h but not as sharply delineated (p=0.132). Again, DRhQ improved performance at 24h for 5xFAD mice (Figure 2B); however, when each sex was evaluated separately, the improvement in NORT performance at 24h was not as clear due to larger noise variance in the vehicle groups. As in the 2h test, there was no interaction between sex and treatment in the 5xFAD mice during the 24h test (p=0.923). There was also no compelling effect of DRhQ treatment in WT mice of either sex in either the 2h (Figure 2A) or 24h (Figure 2B) tests, although the slopes in some groups occasionally appear steep owing to the presence of one or two outlying observations in some groups. In the NORT test there were some animals that were tested but did not engage with either object throughout the duration of the test. Those animals were termed “non-participants” and we found that NORT participation rates did not meaningfully differ by genotype, sex, or DRhQ treatment group, so we believe the treatment effects were not confounded by participation imbalances.

**Figure 2:**
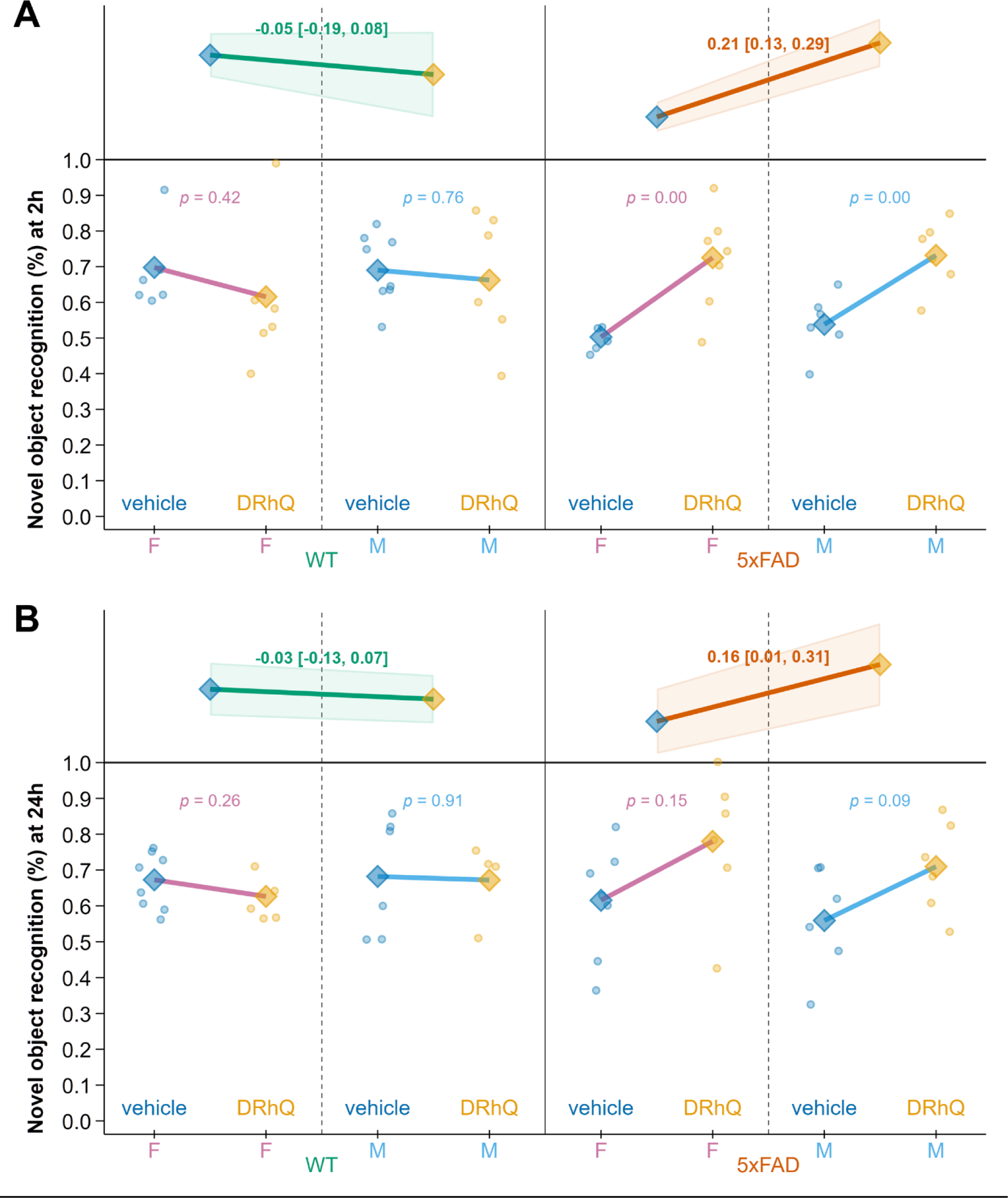
DRhQ improves recognition memory in 5xFAD mice. DRhQ treatment increased time 5xFAD mice spent with the novel object in both the A) 2h and B) 24h test. At 2h, the DRhQ effect was clear in both sexes of 5xFAD animals whereas at 24h the effect of DRhQ was not as clear in either sex individually.

The CFR test evaluates associative memory by recording freezing behavior when mice are reintroduced to the testing chamber 24 hours after receiving a foot shock. Freezing is measured over two five-minute periods to reflect both associative memory retention and extinction. In the first five-minute block (0-5m), there was a trend towards a reduction in freezing in the vehicle-treated 5xFAD animals relative to the vehicle-treated WT animals (p=0.094). In this block there was no salient effect of DRhQ treatment in 5xFAD mice in either sex (Figure 3A) and no interaction between sex and treatment in the 5xFAD mice (p=0.975). There was no visible effect of DRhQ in WT mice. Overall, we were unable to detect the presence or nature of any DRhQ effects in the first five minutes of the CFR test.

**Figure 3:**
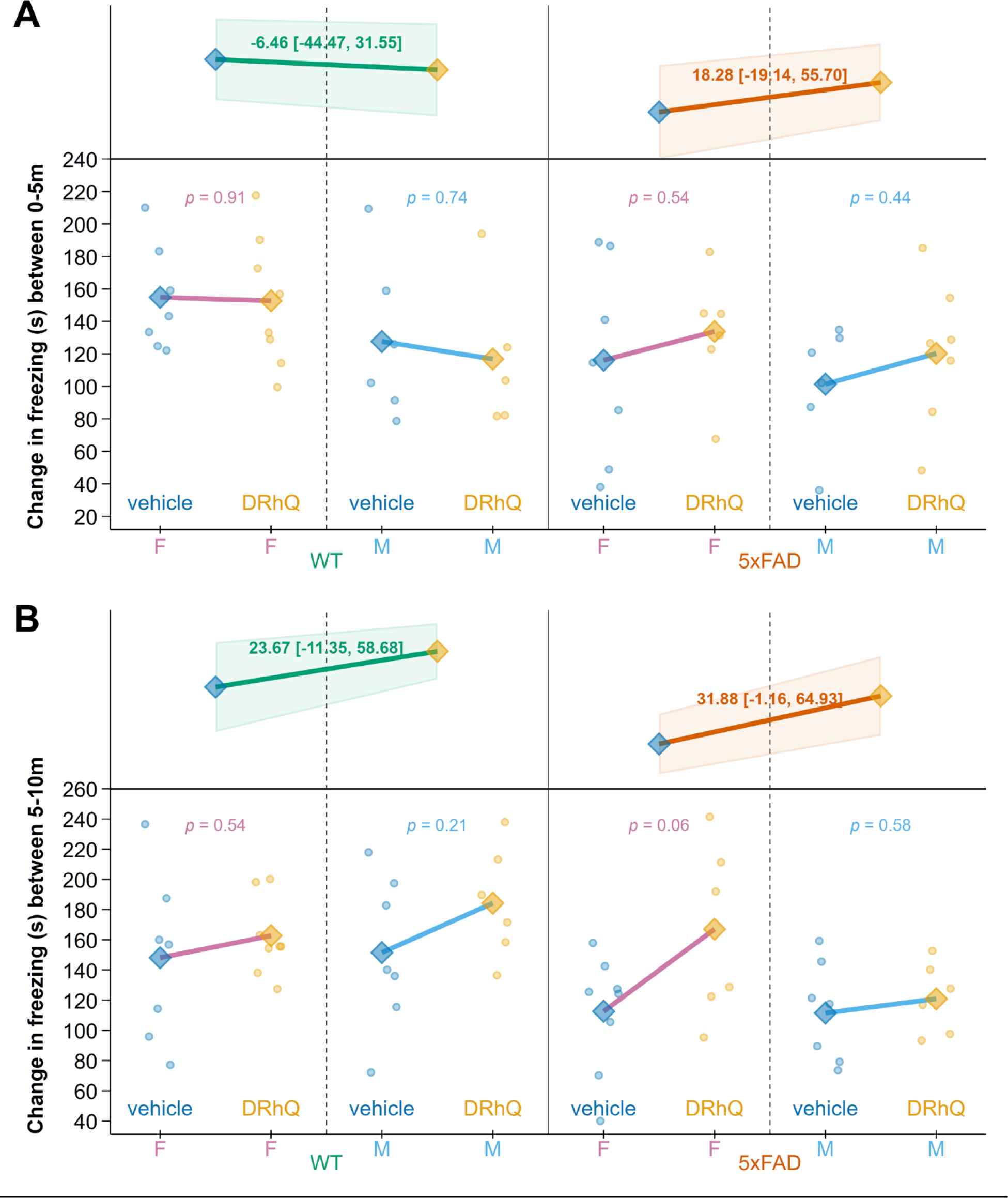
DRhQ treatment does not robustly alter associative memory in 5xFAD mice. A) No unequivocal effects of either genotype or treatment were observed in the first five-minute block of the CFR test. B) In the second five-minute block, there was likewise no clear effect of DRhQ treatment on CFR performance in either the 5xFAD or WT mice.

In the second five-minute block (5-10m), there was a sharper genotype effect with 5xFAD vehicle-treated animals exhibiting impaired performance relative to WT animals in the vehicle treatment group (p=0.039). In this block there was also no clear overall effect of DRhQ in 5xFAD mice (Figure 3B), and no unequivocal interaction between treatment and sex in the 5xFAD mice (p=0.178). Although there was a visible suggestion of an improvement in CFR performance in female 5xFAD mice with DRhQ treatment, it is not possible to say with confidence if that effect is real without the inclusion of more animals (Figure 3B). Overall, it was not clear whether there was a meaningful effect of DRhQ in WT mice in either the first or second block of the CFR test.

### 3.2 DRhQ may reduce Aβ plaque burden in 5xFAD mice only negligibly

Aβ plaque pathology was quantified in 5xFAD mice by the percent area which was determined by a ratio of the Aβ plaque-stained area to the total area of each region (Figure 4A). The pattern of reduction in Aβ plaque burden was consistent in both males and females (more saliently in females) in both brain regions but at such low magnitude in each group that no overall effect of DRhQ on Aβ plaque burden in either the hippocampus or the cortex of 5xFAD mice was resolved at this sample size (Figure 4B and 4C). There was also no clear interaction between sex and treatment in the 5xFAD mice in either region (cortex p=0.362, hippocampus p=0.519). We believe a larger sample size is required to decisively conclude whether a DRhQ effect on Aβ plaque pathology in 5xFAD mice exists, and if so, whether it is truly stronger in females or not.

**Figure 4:**
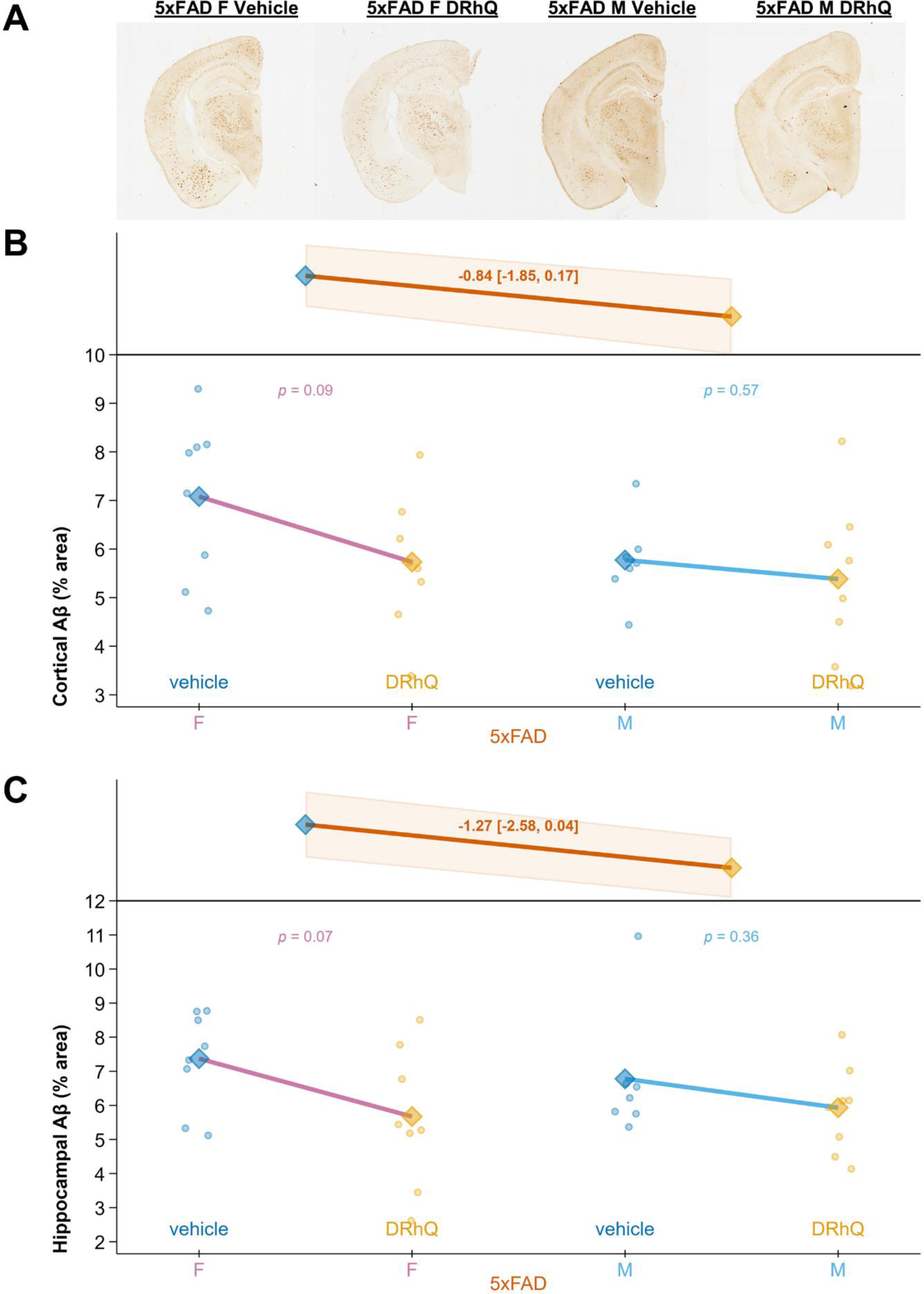
DRhQ treatment does not strongly alter Aβ plaque burden in 5xFAD mice. A) Representative images of Aβ staining in female and male 5xFAD mice. DRhQ treatment had at most a small effect on the overall Aβ plaque burden in both sexes combined in either the B) cortex or C) hippocampus, although in both regions there was a trend towards a reduction in plaque area in all 5xFAD mice, especially in females.

### 3.3 DRhQ improves mitochondrial function in 5xFAD mice

Analysis of mitochondrial function of cortical synaptosomes isolated from WT and 5xFAD mice revealed that vehicle-treated 5xFAD mice had significant impairments relative to WT vehicle-treated mice in basal and maximal respiration (p=0.001 and p<0.001 respectively), as well as spare capacity (p=0.001). A similar suggestion of deficit was seen in the ATP-linked respiration of 5xFAD females compared to WT females, where the 5xFAD mean was visibly lower but with large variance (p=0.151), but not evidently for 5xFAD males (p=0.410), where the means were approximately equal with smaller variance.

Treatment of 5xFAD mice with DRhQ significantly improved basal (Figure 5A), maximal (Figure 5B), and ATP-linked respiration (Figure 5C), as well as spare capacity (Figure 5D). For basal respiration and spare capacity, the DRhQ effect indicated by the overall sex-averaged confidence interval in 5xFAD mice was driven by the changes observed in the female mice whereas for maximal respiration and spare capacity DRhQ elicited improvements in both sexes. For basal respiration there was a trend towards an interaction between sex and treatment in the 5xFAD mice (p=0.121), and indeed the effect of DRhQ in female 5xFAD was visibly more salient than in male 5xFAD, where the effect appeared smaller or perhaps even nonexistent (Figure 5A). There was no interaction between sex and treatment in the 5xFAD mice for maximal respiration (p=0.998) and a significant effect of DRhQ treatment was highly evident in both male and female 5xFAD mice (Figure 5B). There was a fairly sharp interaction between sex and treatment in 5xFAD mice for ATP-linked respiration (p=0.029), with only female 5xFAD mice showing any improvement with DRhQ treatment (Figure 5C). A trend towards an interaction between sex and treatment in 5xFAD mice was also evident for spare capacity (p=0.150) and although a similar and highly salient improvement in spare capacity was seen with DRhQ treatment in both sexes of 5xFAD mice (Figure 5D in the male 5xFAD mice it may be due to the presence of two anomalously low outlying values (much lower than any other observations) in the 5xFAD male group. There was no consistent or compelling effect of DRhQ treatment in WT mice for any metric of cortical mitochondrial bioenergetics.

**Figure 5:**
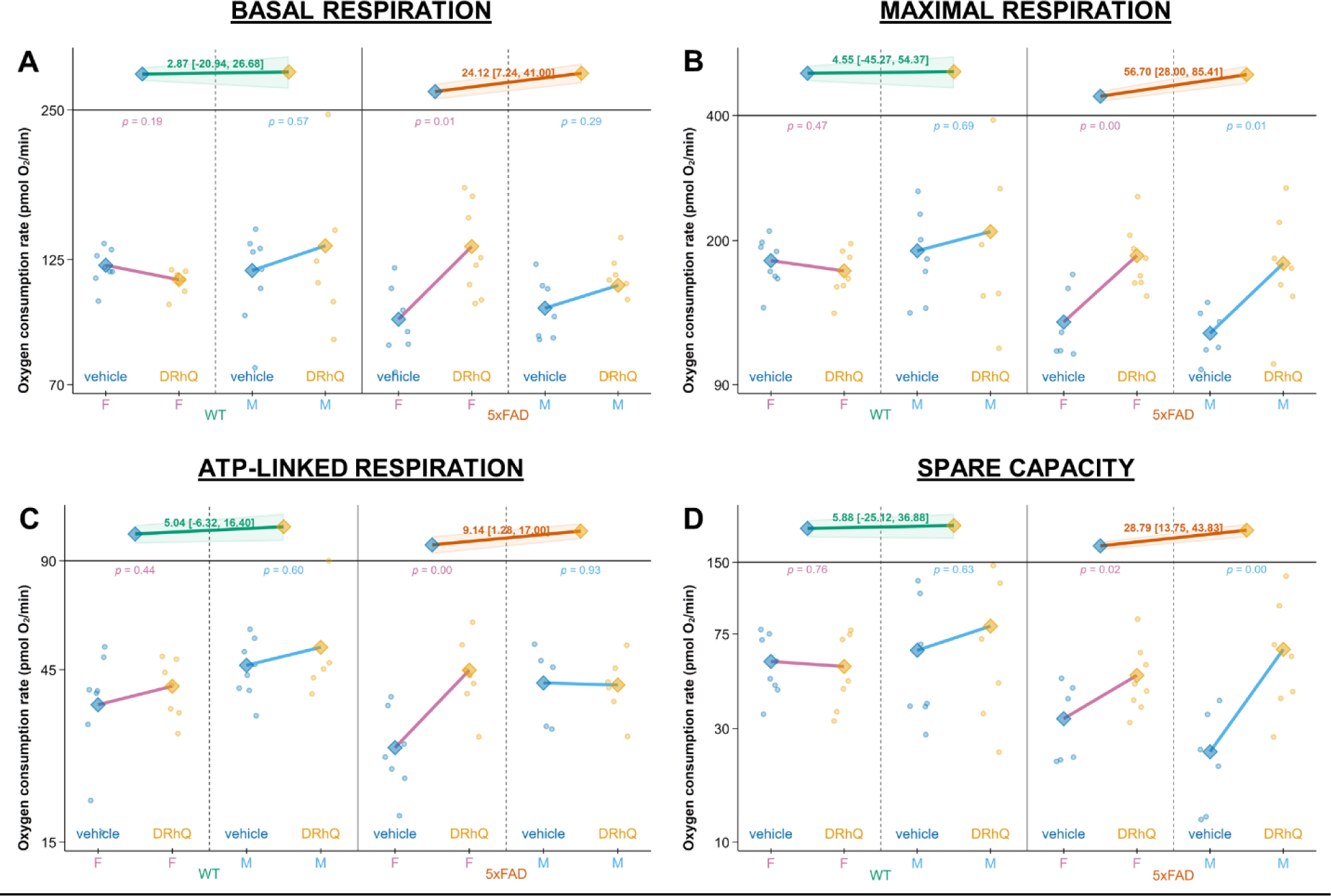
DRhQ improves mitochondrial respiration in 5xFAD mice. DRhQ treatment increased A) basal, B) maximal, and C) ATP-linked mitochondrial respiration as well as D) spare capacity in cortical synaptosomes isolated from 5xFAD mice. The DRhQ -induced changes in maximal respiration and spare capacity were apparent in both sexes while the effect of treatment on basal and ATP-linked respiration was visible only in female 5xFAD mice. DRhQ did not meaningfully affect any of these metrics in WT mice.

**Figure 6:**
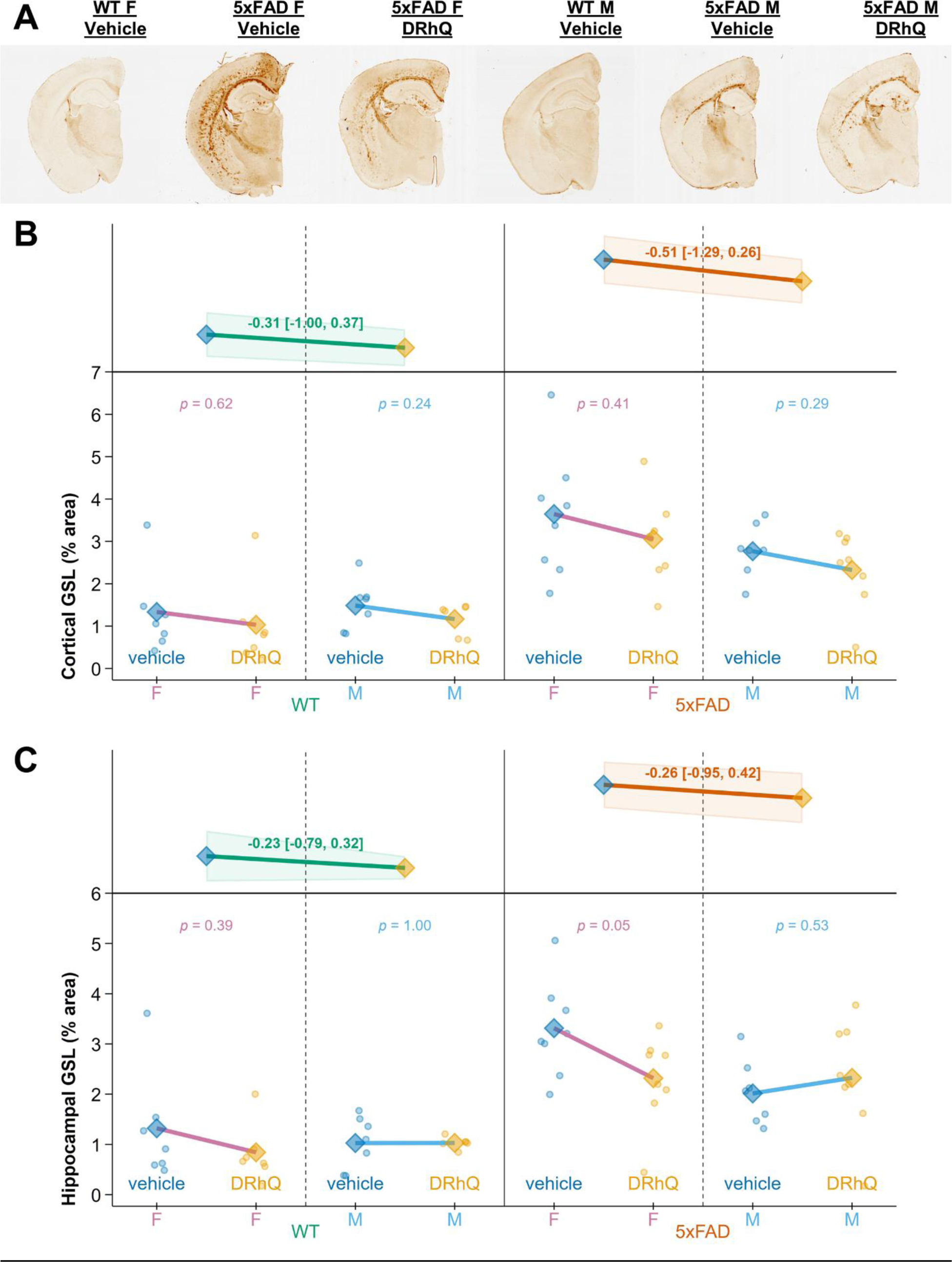
DRhQ treatment did not consistently alter microglial activation in 5xFAD mice. A) Example images from female and male mice. B) There was no clear change in GSL staining in the cortex of either 5xFAD or WT mice. C) There was similarly no visible effect of DRhQ treatment in the hippocampus of 5xFAD mice when both sexes were evaluated together; however, in 5xFAD females the possibility of a reduction was observed in the DRhQ-treated mice.

The expression of genes encoding mitochondrial enzymes in the ETC was also evaluated in synaptosomes isolated from the cortex of treated animals. Reduced cortical expression of the ETC gene Mt-ND1 was seen in 5xFAD mice compared to WT mice and a similar, but less clear, trend was observed in Mt-CO1 and Mt-ATP6 (Table 1). However, a combined index (linear combination) of the three gene expression values showed a sharper effect of genotype (p=0.002; data not shown) than any of the genes analyzed alone. However, DRhQ treatment did not appear to alter the expression of any of these genes in either 5xFAD or WT mice.

**Table 1:**
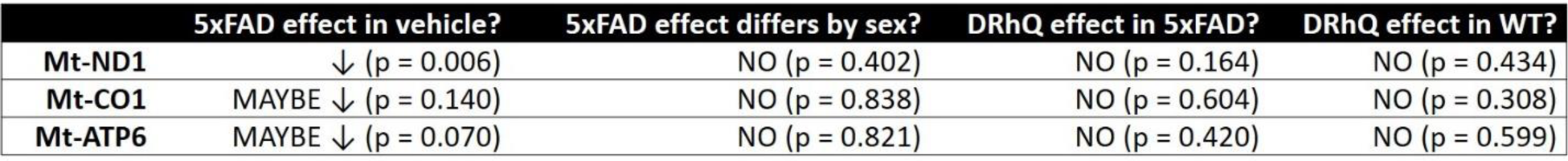
DRhQ treatment did not change cortical mitochondrial gene expression. Interpretations of effects and p-values for gene expression changes.

|  | 5xFAD effect in vehicle? | 5xFAD effect differs by sex? | DRhQ effect in 5xFAD? | DRhQ effect in WT? |
| --- | --- | --- | --- | --- |
| **Mt-ND1** | ↓ (p = 0.006) | NO (p = 0.402) | NO (p = 0.164) | NO (p = 0.434) |
| **Mt-CO1** | MAYBE ↓ (p = 0.140) | NO (p = 0.838) | NO (p = 0.604) | NO (p = 0.308) |
| **Mt-ATP6** | MAYBE ↓ (p = 0.070) | NO (p = 0.821) | NO (p = 0.420) | NO (p = 0.599) |

### 3.4 DRhQ does not robustly affect markers of neuroinflammation in 5xFAD mice

GSL (*Griffonia simplicifolia* Lectin I) staining was measured as a marker of microglial activation immunohistochemically (Figure 5A). Increased cortical and hippocampal microglial activation was observed in 5xFAD mice of both sexes compared to WT animals (p<0.001 for both the cortex and hippocampus). DRhQ treatment did not evoke a clear or consistent effect on GSL staining in 5xFAD mice. In the cortex there was no evidence of interaction between sex and treatment in the 5xFAD mice (p=0.857) and a hint of s negative trend in each sex. However, in the hippocampus there was a suggestion of interaction (p=0.063) because the female and male trends were in opposite directions, suggesting the possibility of sex-specific effects on microglial activation. There was some evidence of a reduction in GSL staining in the hippocampus of female 5xFAD mice treated with DRhQ (Figure 5C), but this outcome too requires a larger sample size before a differential DRhQ effect due to genotype and sex could reasonably be asserted. Overall, the empirical trends in the effects of DRhQ on GSL staining observed in this study lack resolution to draw firm conclusions.

Similarly, although a subset of the proinflammatory markers assessed were increased in 5xFAD relative to WT in vehicle-treated mice, DRhQ did not alter the expression of any inflammatory genes in the cortex of treated 5xFAD animals (Table 2). Interestingly, DRhQ treatment did elicit a reduction in TNFα expression, but this was only evident in WT mice and not observed for any other genes evaluated and the trend was driven by just a few unusually influential observations.

**Table 2:** DRhQ treatment did not alter cortical expression of inflammatory genes in 5xFAD mice. Interpretations of effects and p-values for gene expression changes.

|  | 5xFAD effect in vehicle? | 5xFAD effect differs by sex? | DRhQ effect in 5xFAD? | DRhQ effect in WT? |
| --- | --- | --- | --- | --- |
| IL6 | ↑ (p = 0.015) | NO (p = 0.140) | NO (p = 0.622) | NO (p = 0.246) |
| TNFα | ↑ (p = 0.006) | NO (p = 0.145) | NO (p = 0.458) | ↓ (p = 0.011) |
| TREM2 | ↑ (p = 0.003) | NO (p = 0.505) | NO (p = 0.886) | NO (p = 0.326) |
| C3AR1 | ↑ (p < 0.001) | NO (p = 0.158) | NO (p = 0.760) | NO (p = 0.267) |
| CX3CR1 | ↑ (p = 0.010) | NO (p = 0.241) | NO (p = 0.469) | NO (p = 0.682) |
| CX3CL1 | NO (p = 0.888) | NO (p = 0.452) | NO (p = 0.754) | NO (p = 0.798) |
| RAGE | NO (p = 0.686) | NO (p = 0.353) | NO (p = 0.816) | NO (p = 0.960) |
| CCR6 | NO (p = 0.841) | NO (p = 0.111) | NO (p = 0.755) | NO (p = 0.791) |
| CD36 | NO (p = 0.898) | NO (p = 0.111) | NO (p = 0.529) | NO (p = 0.247) |
| CD74 | NO (p = 0.342) | MAYBE (p = 0.071) | NO (p = 0.262) | NO (p = 0.184) |

## 4. Discussion

This study sought to determine potential therapeutic impacts of DRhQ treatment on the 5xFAD model of AD. The data collected shows promising signs of potentially mitigating effects of the novel treatment on AD pathology and its downstream consequences.

DRhQ treatment significantly improved recognition memory in 5xFAD mice. Although to our knowledge neither DRhQ nor its predecessor DRɑ1 have been evaluated for the effects on cognition, MIF inhibition by other means has similarly improved cognitive function in an AD mouse model. In a streptozotocin-induced model of AD the MIF inhibitory compound ISO-1 improved associative memory [29]. Interestingly, in our study we did not observe a clear effect of DRhQ on associative memory. This could reflect regional specificity of the action of DRhQ, but that seems unlikely given the reported beneficial effects of DRhQ and DRɑ1 in a wide array of neurodegenerative conditions mediated by diverse brain regions [20]. Another possibility is that our relatively small sample size prevented the detection of a significant treatment effect. This is plausible since we also did not observe a significant genotype effect in 5xFAD mice, which is not consistent with what has been widely reported by our group and others [30–34]. Future studies with more animals are therefore needed to clarify the ambiguous results of MIF-inhibition on associative memory seen here.

Administration of DRhQ also did not strongly affect Aβ plaques in 5xFAD mice. This was somewhat surprising given the association between MIF and Aβ that has been documented in both preclinical and clinical settings. Elevated MIF expression has been observed in the plasma and CNS of AD patients [13, 14] and in APP23 transgenic mice, MIF was seen to co-localize with Aβ plaques [19]. We therefore expected that treatment of 5xFAD mice with the MIF inhibitor DRhQ would result in decreased Aβ plaques. We did in fact observe mild but consistent trends suggestive of reduced plaques, but the size of effect was too small and the resolution provided by our limited sample size too weak to resolve the trends with any clarity. Our data in female 5xFAD mice provides the strongest hint of the possibility of a trend toward a reduction in Aβ plaques in the cortex and hippocampus in line with this idea, although as mentioned, a future study with many additional animals would be necessary to determine those effects with certainty.

We also found that DRhQ administration improved cortical mitochondrial function in 5xFAD mice. Generally, this effect was more pronounced in the female 5xFAD animals. This may be due to the fact that male 5xFAD mice had less plaque pathology than female 5xFAD mice, a phenomenon that we and other groups have previously reported in 5xFAD mice [31]. In fact, in our study we observed a similar trend towards increased plaque burden in the cortex of female mice (p=0.055). If the plaque burden in male 5xFAD mice is lower, then it would also be expected that the deleterious consequences of Aβ on mitochondrial function would likewise be more subtle. There was no effect of DRhQ on cortical mitochondrial gene expression in either sex. However, this suggests that the impact of DRhQ, in female 5xFAD mice at least, was on mitochondrial function itself rather than through changes in overall mitochondrial content or biogenesis. While effects of MIF silencing or inhibition on mitochondrial dynamics and maintenance of mitochondrial membrane potential have been reported in cancer cells [35, 36], to our knowledge this is the first report of effects of inhibiting MIF-CD74 signaling on brain mitochondria in the context of Aβ accumulation.

Surprisingly, we did not observe a robust effect of DRhQ on the neuroinflammatory markers assessed in this study. We did not see any changes in inflammatory gene expression in the cortex of DRhQ-treated 5xFAD mice. These findings are not consistent with previously reported beneficial effects of DRα1 on neuroinflammation observed in models of multiple sclerosis, stroke, traumatic brain injury and methamphetamine addiction [20, 22–25]. This discrepancy may be because the effects of DRhQ are largely specific to microglia, and perhaps by extracting RNA from whole tissue homogenates rather than isolated microglial cells, we were not able to detect a treatment effect. In fact, when we looked at microglial activation via GSL staining there was some evidence of an effect of DRhQ in the hippocampus of female mice with a possible interaction between sex and treatment effect in 5xFAD mice (p=0.063) and an effect of treatment in female animals (p=0.05). The fact that a suggestion of a treatment effect is more apparent in female 5xFAD mice rather than male mice again could be related to the differential Aβ plaque burden reported between the sexes.

Taken together, the results from this study suggest that DRhQ has promise as an AD therapeutic agent. Further studies with more animals are needed to elucidate the entire spectrum of its cellular effects, optimize dosing, and to more fully understand potential sex differences in response to the drug.

## Acknowledgments

This work was funded by a pilot grant from the Oregon Alzheimer’s Disease Research Center (NIA P30AG066518, PI: Jeffrey Kaye).

---

**Supplementary Table 1.**
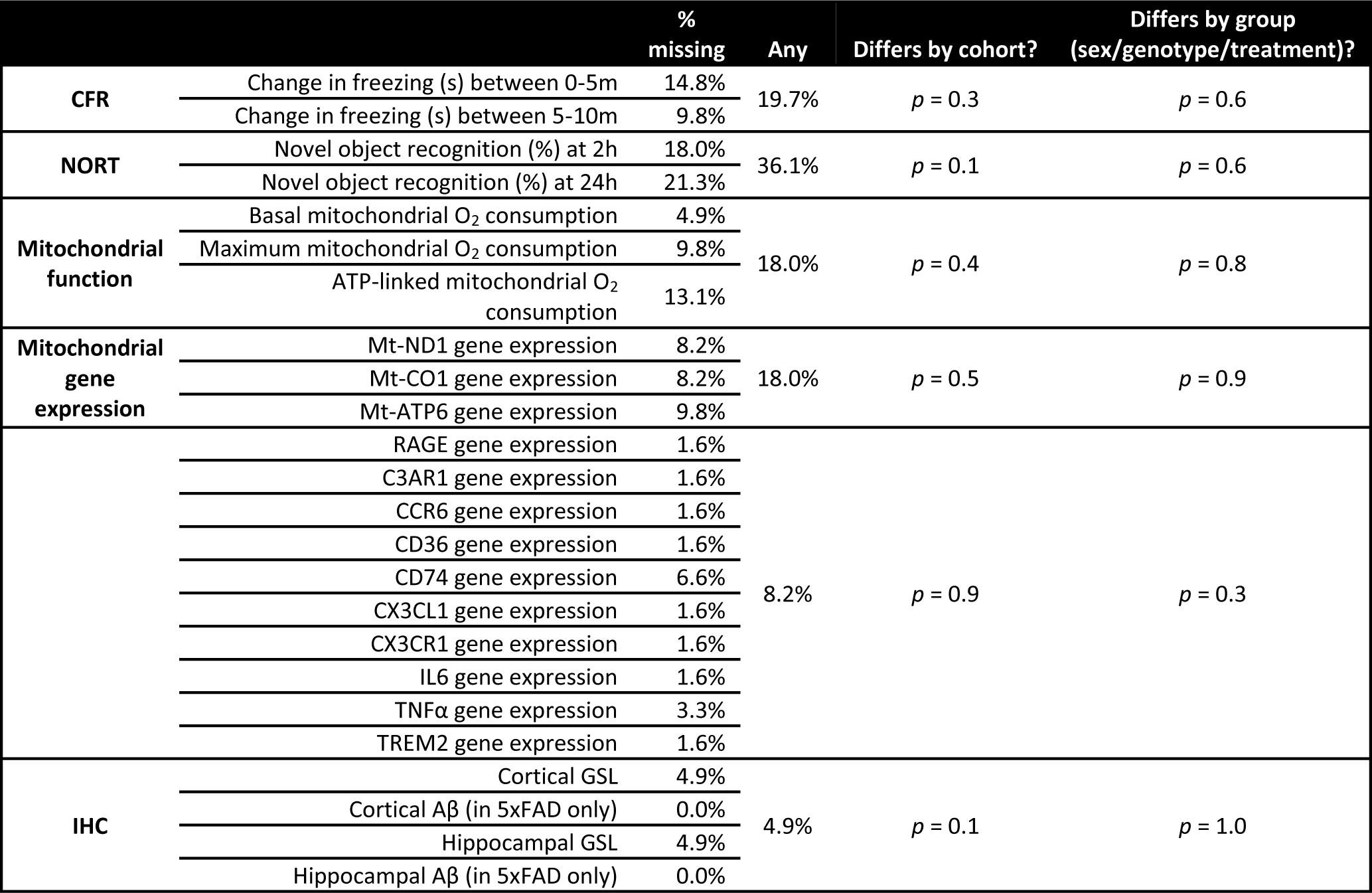

